# A new Dicistrovirus from soldier fly *Inopus flavus* (James) (Diptera: Stratiomyidae), a pest of sugarcane

**DOI:** 10.1101/2020.12.24.411611

**Authors:** Angelique K. Asselin, Kayvan Etabari, Michael J. Furlong, Karyn N. Johnson

## Abstract

The native Australian soldier flies, *Inopus* spp. (Diptera: Stratiomyidae), are agricultural pests of economic importance to the sugarcane industry. While adult soldier flies do not feed on sugarcane, larvae spend one to two-years underground feeding on roots, causing mechanical and systemic damage to crops (*Saccharum officinarum L.*) that impacts yield. Current measures of pest control commonly target above ground pests and are ineffective against solider fly larvae, highlighting the importance of novel control methods. A screen of the salivary gland transcriptome of *Inopus flavus* (James) revealed the presence of viral RNA belonging to a potentially novel member of the *Dicistroviridae* family. Viruses from this family have been found naturally infecting insects from a range of taxonomic groups and they often cause pathogenesis in their hosts. To characterise the genetic and physical properties of the new virus, the positive RNA genome was analysed using a combination of sequencing approaches. The virus genome is organised similarly to members of the *Dicistroviridae* with two open reading frames (ORF) the first encoding non-structural proteins and the second encoding structural proteins. The genome includes two potential internal ribosomal entry sites (IRES) one within the 5’ UTR and the other in the intergenic region (IGR). Based on the amino acid sequences of the non-structural and structural polyproteins encoded by the two ORF soldier fly virus groups within the dicistrovirus family. Virus particles purified from infected larvae and visualised by electron microscopy are icosahedral, non-enveloped, and 30 nm in diameter. The genetic and physical characteristics of this novel soldier fly virus are consistent with it being a member of the *Dicistroviridae*.

---

Australian soldier flies, *Inopus* spp. (Diptera: Stratiomyidae), are endemic to eastern Australia and southern Papa New Guinea and *Inopus rubriceps* has been introduced into the North Island of New Zealand and California, USA [1, 2]. *Inopus* spp. are pests of economic importance across both their native and introduced ranges where thay attack sugarcane, pasture, horticultural and fodder crops [1]. The subterranean larvae live for 1-2 years and their sucking mode of feeding on roots causes systemic damage to above ground parts of affected plants [3–7]. In sugarcane, feeding presents as inhibited germination and retarded shoot development [8, 9] and it can result in significant yield loss. Although the mechanism of systemic damage remains unclear, larval homogenates reduce sugarcane sett germination [10], and feeding on sugarcane roots upregulates venom proteins in larvae, suggesting these proteins could be involved in damage to host tissue [11, 12]. In contrast, adult soldier flies do not cause plant damage and they are ephemeral, living for 3 to 10 days depending on location, during which time they mate and lay eggs close to the soil surface near host plants [3, 13, Reviewed in 14].

Currently there is no effective control for soldier flies. Management approaches have included targeting of adults with pesticides in the short temporal window after emergence [15] but this is ineffective. Control of larvae is also difficult. Insecticide based approaches have failed due to variable efficacy of compounds and the need to integrate applications with specific growing practices [14, 16–18]. The cost, ineffectiveness and negative non-target impacts of pesticides, the lack of effective arthropod natural enemies [19] and poor control provides by previous management practices [Reviewed in 14, 20] have prompted investigations for alternative control measures.

The broad range of hosts, pathology and effects on host behaviour make dicistroviruses appealing as biopesticides for control of agricultural pests. Members of *Dicistroviridae* infect a variety of arthropod hosts. They principally attack insects and they can have broad host ranges, for example Cricket paralysis virus (CrPV) infects multiple insect orders [21–28]. Arthropod hosts of medical and agricultural importance are infected and viruses attacking agriculturally important hosts include CrPV [26, 27], Aphid lethal paralysis virus (ALPV) [29], Rhopalosiphum padi virus (RhPV) [30, 31], Plautia stali intestine virus (PSIV) [32], Himetobi P virus (HiPV) [33], Homalodisca coagulate virus-1 (HoCV-1) [34] and Solenopsis invicta virus-*1* (SINV-1) [35]. Infections vary from asymptomatic and chronic [23, 36–39] to acute and highly pathogenic, resulting in rapid disease progression and host death [26, 29, 40, 41]. Chronic infections, such as those seen with *Drosophila C virus* (DCV) [39], can be asymptomatic or mildly pathogenic but stress or secondary infections may trigger increased pathogenicity and more rapid disease progression [23, 37, 42–44]. More symptomatic infections commonly present with paralysis [26, 29, 40, 41], reduced longevity or fecundity [31, 39], reduced larval and pupal longevity [45] or changes in behaviour [46]. The combination of host range, pathology and resulting effects on host interactions with enviroment make dicistroviruses promising for biological control [43].

The *Dicistroviridae* family is characterised by single stranded, positive sense, non-partite, RNA genomes contained within non-enveloped icosahedral virions approximately 30 nm in diameter [47]. The 5’ end of the viral genome is covalently linked to a viral protein genome-linked (VpG) and ranges between 8-10kb, encoding two open reading frames (ORF) and poly-adenylated. ORF1 is located at the 5’ end and encodes a non-structural polyprotein comprised of an RNA dependent RNA polymerase (RdRp), protease, helicase and VpG, ORF2 is separated from ORF1 by an intergenic region (IGR) and encodes a structural capsid polyprotein producing four separate structural proteins VP1, VP2, VP3, and VP4 [47]. Upon translation the polyproteins’ are processed into individual components by a virally encoded proteinase [47]. In some members, there is a third overlapping +1 frame ORFx within ORF2 [48–50] that appears to be important for CrPV infection [51]. Dicistroviruses employ a non-canonical method of translation through the use of internal ribosome entry sites (IRES) at both the 5’UTR and IGR for translation of ORF1 and ORF2 respectively. The structure and function of the 5’UTR IRES of dicistroviruses has not been well characterised. Despite an apparent lack of conserved structures across the family, they appear to function as IRESes *in vitro* and require the recruitment of initiation factors [52–54]. The 5’UTR IRES of CrPV, has been elucidated with the structure and function of a group III IRES. Meanwhile the IGR IRES is group IV, the secondary structure of which allows recruiting of 80S ribosomal subunits directly to initiate translation. The IGR IRES of dicistroviruses have been extensively studied and can be categorised as either Type I or II based on conservation of distinct structural elements not necessarily sequence conservation. Triatoviruses and cripaviruses seem to have type I while aparaviruses have type II. The main domains of the IGR IRES include stem loops and bulge regions (L1.1, SLIV, SLV) and pseudoknots (PK I, II, and III). While Type II have an extra stem-loop domain SLIII within PKI and a bigger L1.1 domain the two IRES Types function similarly in initiating translation [55–58]. Here we describe a new soldier fly virus which may prove to be a pathogen of these pests and which could be exploited for their control.

Soldier fly larvae were collected from the stools of infested sugarcane (*Saccharum officinarum L.*) fields near Hay Point, Mackay, Queensland Australia (21°18′5″S,149°14′7″E) as described [12] in February and March 2019, prior to annual emergence of adults. RNA, purified from larvae as described previously and used to generate RNAseq data [12], was also used to synthesise viral cDNA using specific primers **(Table 1)** with SuperScript® Double-Stranded (Invitrogen by Life technologies) as per manufacturers instructions. Further rounds of cDNA were synthesised using a the SMARTer RACE 5’/3’ Kit (Clontech Laboratories, Takara Bio) as per manufacturers instructions. cDNA was cloned into the pGEM®-T Easy Vector (Promega) per manufacturers protocol and transformed into competent XL1 *Escherichia coli* cells by heat shock. Plasmid DNA was purified from positive clones using a ZR Plasmid Miniprep – Classic kit (Zymo Research) following manufacturer’s instructions and 600-1000 ng of pure DNA in UltraPureTM water (Invitrogen)) were sent to the Australian Genome Research Facility for Sanger sequencing using sequencing primers **(Table 1).** A consensus of the novel soldier fly virus genome was derived using RNAseq data (GSE127658) and Sanger sequencing of cloned cDNA. The 9 762 nt RNA genome (excluding poly(A) tail) was compiled using RNAseq data accession number MW357714 (Figure 1A). With an average coverage of 402 728 reads per nucleotide, the consensus represents the most prevalent sequence in the RNAseq library, with only 26 single nucleotide variants (SNV), 3 multi nucleotide variants and one deletion with Phred scores above 29 and frequencies above 10% (Figure 1B and 1C). The consensus was confirmed by Sanger sequencing with 93.35% homology over 9 206 nt excluding the 5‘ end. Considering the RNA template used for sequencing was derived from a pool of soldier fly larvae the variation between sequences and in the library likely reflects the diversity of the viral population, which is expected as RNA viruses largely exist as quasispecies [59].

**Table 1:**
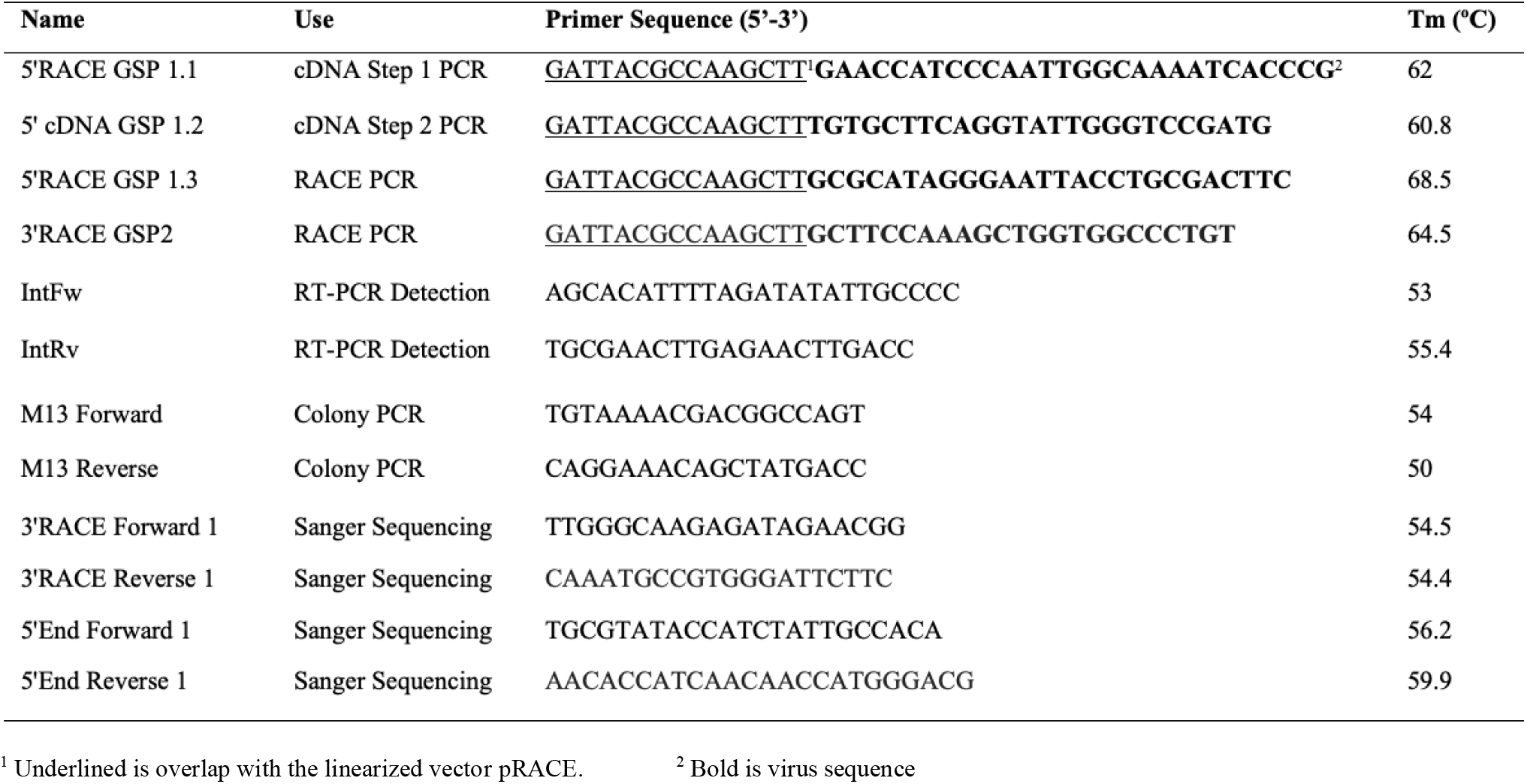
Nucleotide sequence of primers used in this publication.

**Figure 1:**
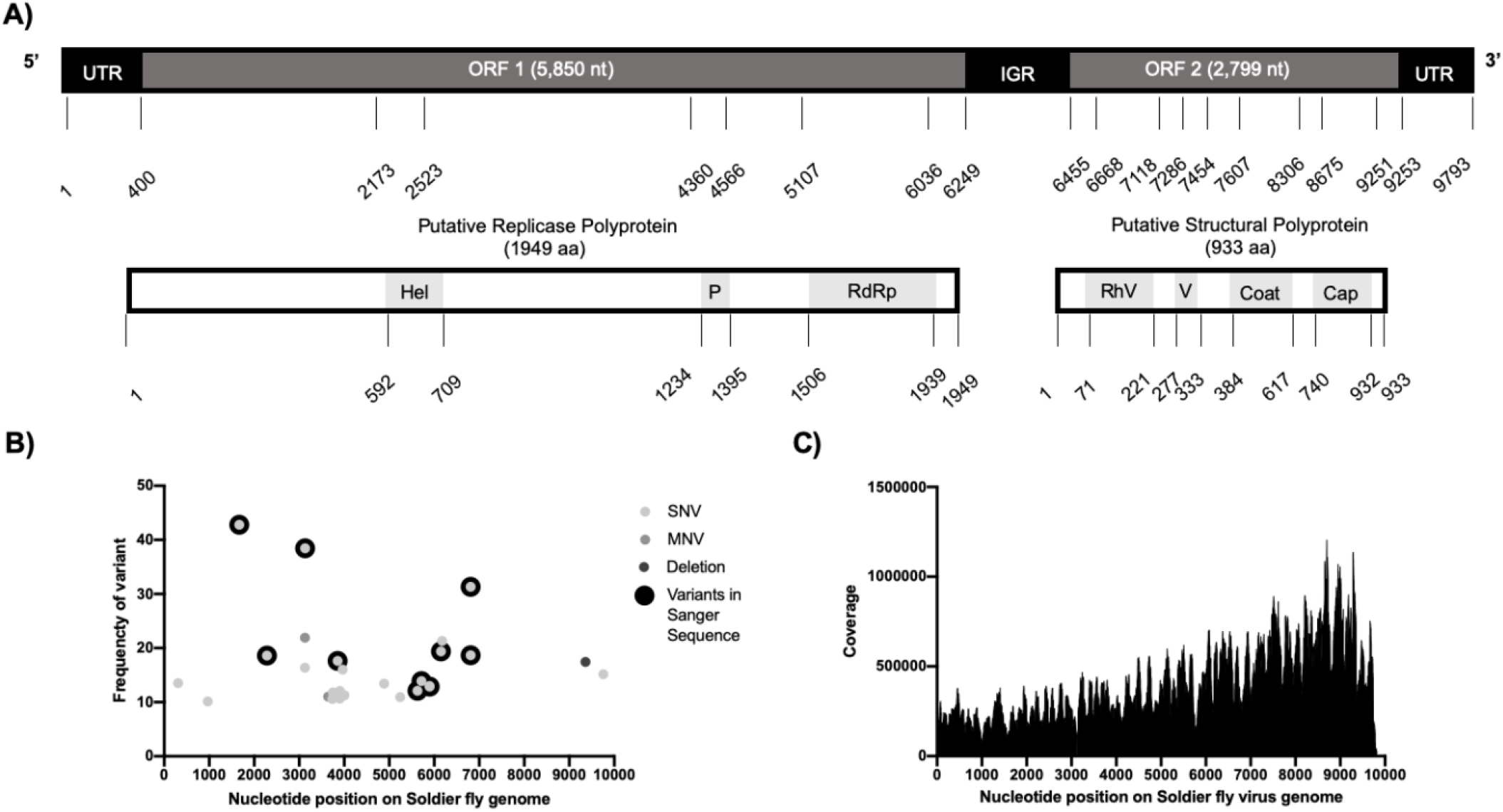
**A)** Schematic of the organization of the new soldier fly virus genome. The conserved structural domains of the two open reading frames (ORF) predicted by HMMER 3.1b1 (May 2013) in CLC Main Workbench 8.1 search of the Pfam-A v32 database. Domains for ORF1 include Hel = RNA helicase (PF00910.22, E-value: 1.30e-17), Peptidase = Peptidase C3G (PF12381.8, E-Value: 1.10e-4), RdRp = RNA dependent RNA polymerase (PF00680.20, E-Value 2.20e-48) and for ORF 2 RhV = picornavirus capsid protein (PF00073.20, E-Value: 2.90e-12) V = Cricket paralysis virus VP4 (PF11492.8, E-Value: 1.8e-8), Coat = Calicivirus coat protein (PF00915.20, E-Value: 8.60e-5), Cap = CrPV capsid protein like (PF08762.10, E-Value: 3.3e-17). **B)** Map of single nucleotide variants (SNV) multi-nucleotide variants (MNV) and deletions present in the RNAseq library with a frequency above 10% and with quality Phred scores above 29. There were also high frequency variants found in the Sanger Sequence consensus. Variants appear across the genome with no apparent regions of high variation. **C)** RNASeq read density plotted along the solider fly virus genome. Y-axis the number of reads mapped to a particular nucleotide of the soldier fly virus genome X-axis. There is higher coverage closer to the 3’ end that contains a poly(A) tail.

Comparison of the assembled genome to the NCBI-nucleotide database identified sequence similarity with dicistroviruses. The genome organisation of this putative new virus is two ORFs separated by an IGR with both a 5’ and 3’ UTR and poly(A) tail (Figure 1A). The first ORF predicted by CLC Main Workbench with an AUG (Met) start codon and translated using standard genetic code translation encodes a putative non-structural polyprotein containing three motifs an RNA helicase (PF00910.22, E-value: 1.30e-17), Peptidase C3G (PF12381.8, E-Value: 1.10e-4) and RNA dependent RNA polymerase (PF00680.20, E-Value 2.20e-48) (HMMER 3.1b1 (May 2013) in CLC Main Workbench 8.1 using the Pfam-A v32 database) (Figure 1A).

Translation of ORF2 in dicistroviruses is not intiated by a canonical methionine codon [57, 58], commonly it is GCU (Ala) [Reviewed in 60]. It was not possible to robustly identify conserved motifs of dicistroviruses IRESs in the IGR of the soldier fly virus. Without prediction of IRES structures it was not possible to predict the start codon of reading frame and analysis of ORF2 was conducted from the first in frame methionine. The amino acid sequence encoded by the second ORF contains motifs that match structural polyprotein sequence of other dicistroviruses, including picornavirus capdis protein (RhV, PF00073.20, E-Value: 2.90e-12), Cricket paralysis virus VP4 (Dicistro VP4, PF11492.8, E-Value: 1.8e-8), Calicivirus coat protein (Calic_coat, PF00915.20, E-Value: 8.60e-5), CrPV capsid protein like (CrPV, PF08762.10, E-Value: 3.3e-17) (Pfam-A v32 database search on CLC Main Workbench 8) (Figure 1A). The organisation of the RNA genome of soldier fly virus and its sequence homology with other dicistroviruses supports its classification as a member of this family.

A multiple-sequence alignment of the protein sequence of ORF1 and ORF2 of the 15 accepted members of the dicistrovirus family and tentative members from metagenomic data was created using the following parameters, Gap open cost = 10, Gap extension cost = 1.0, End gap cost = As any other (CLC Main Workbench 8.1). The alignment was used to create an un-rooted maximum likelihood phylogeny using a neighbor joining construction method with a WAG protein substitution rate (CLC Main Workbench 8.1, Figure 2).

**Figure 2:**
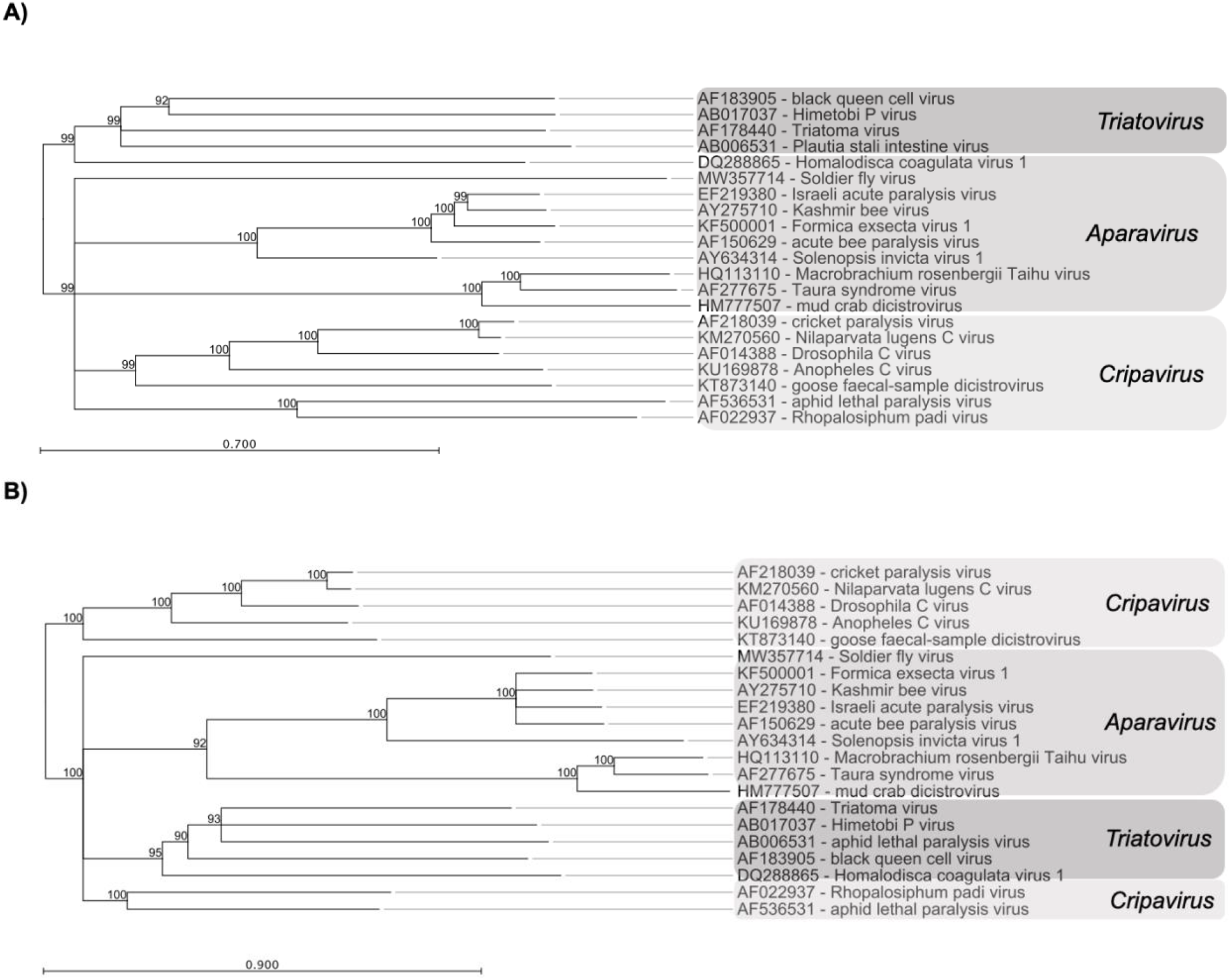
A multiple-sequence alignment of the protein sequence of **A)** ORF1 and **B)** ORF2 of the 15 accepted members of the Dicistrovirus family (black) and tentative members from metagenomic data (*) was used to create an un-rooted maximum likelihood phylogeny using a neighbor joining construction method with a WAG protein substitution rate. One thousand bootstraps were performed and only branches with bootstrap values greater than 90% are shown. Viruses used for the alignment, AB006531 - Plautia stali intestine virus, AB017037 - Himetobi P virus, AF014388 - Drosophila C virus, AF022937 - Rhopalosiphum padi virus, AF150629 - acute bee paralysis virus, AF178440 - Triatoma virus, AF183905 - black queen cell virus, AF218039 - cricket paralysis virus, AF277675 - Taura syndrome virus, AF536531 - aphid lethal paralysis virus, AY275710 - Kashmir bee virus, AY634314 - Solenopsis invicta virus, DQ288865 - Homalodisca coagulata virus 1, EF219380 - Israeli acute paralysis virus, HM777507 - mud crab dicistrovirus, HQ113110 - Macrobrachium rosenbergii Taihu virus, KF500001 - Formica exsecta virus 1, KM270560 - Nilaparvata lugens C virus, KT873140 - goose faecal-sample dicistrovirus.

Dicistroviruses are grouped into three putative genera, *Aparavirus, Cripavirus* and *Triatovirus* depending on the phylogenetic distance of ORF polyprotein sequences and IGR IRES structure. It should be noted however that *Aparavirus* often separates into two divergent clades, one containing viruses that infect insects and the other viruses that infect crustacea [47]. As mentioned previously the IRES structure of soldier fly virus could not be predicted, but the the phylogenies of ORF1 and ORF2 groups soldier fly virus within the dicistrovirus family (Figure 2).

Pools of 8-10 soldier fly larvae and adults were screened for the presence of virus using RT-PCR and all samples were positive (data not show). Positive pools of larvae were homogenised in 500 μL phenylthiourea (PTU) saturated phosphate buffered saline (PBS, pH 7.4) to prevent melanisation. Virus was purified from positive larvae samples by ultracentrifugation on a 10-40% sucrose-density gradient in phosphate buffer (PB), essentially as described for other dicistroviruses [61, 62]. Briefly, virus was diluted with PB and centrifuged on a 10 % sucrose cushion (27,000 rpm, 3 h, 12°C). Supernatant was removed and the pellet was resuspended by vortexing and storage at 4 °C overnight in 250 μL of PB. The resuspended pellet was transferred to a microcentrifuge tube and centrifuged in a sigma 3-16kl benchtop centrifuge for 2-5 minutes at 14,000 rpm to remove any remaining particulate matter. The supernatant was then centrifuged through a 10-40% sucrose gradient at 12°C and 27,000 rpm for 2 hours. Where not specified, centrifuge steps were carried out in in SW41 tubes (Beckmann Coulter, Life Science) using a SW41 swing rotor in an Optima L-80XP Ultracentrifuge (Beckmann Coulter, Life Science). One mL fractions of the gradient were collected, and viral protein was detected by 12% SDS-PAGE gel, stained with Coomassie Brilliant Blue G-250. RT-PCR as previously described confirmed the identity of the purified virus as novel soldier fly virus (data not shown). 12% SDS-PAGE of the purified virus revealed two protein bands approximately 32 and 31 kDa in size and one at 50 kDa (Figure 3A), likely corresponding to the capsid proteins. We we were unable to detect protein bands of lower molecular weight. While dicistroviruses have four capsid proteins (VP1, VP2, VP3 and VP4), sometimes only two dominant bands resolve [63, 64]. Purified virus samples were sent to the Centre for Microscopy and Microanalysis (CMM), The University of Queensland, for transmission electron microscopy imaging. Samples were negatively stained in 2% uranyl acetate and visualized with by electron microscopy. Non-enveloped icosahedral viral particles of an approximate 30 nm-diameter can be seen (Figure 3B), consistent with a member of *Dicistroviridae*.

**Figure 3:**
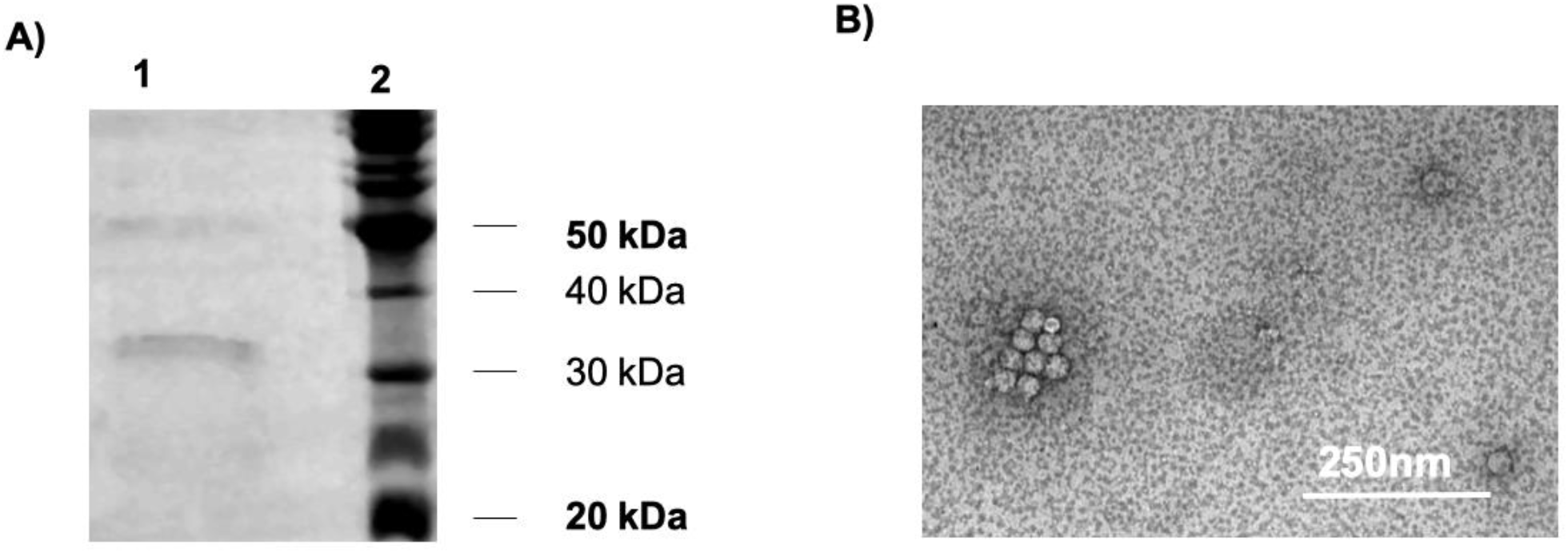
Visualisation of soldier fly virus particles purified from *Inopus flavus* larvae. **A)** A Coomassie brilliant blue stained 12% SDS-PAGE. Lane 1: purified soldier fly virus showing two protein bands approximately 30 kDa and one at 50 kDa likely the viral capsid proteins Lane 2: Benchmark™ Unstained Protein Ladder. **B)** Transmission electron microscopy of Solider fly virus particles diluted 1/5 in phosphate buffer (pH 7.2) and negatively stained with 2% uranyl acetate on a Cu 400 home made glow discharged grid. Virus particles appear either as isolated particles or aggregated. The shape of the viral particles is icosahedral and they are approximately 30nm-diameter.

## Summary

We identified and purified a novel ss +RNA virus from Australian *Inopus* spp. larvae that has genetic and physical properties similar to members of the *Dicistroviridae* family. Dicistrovirus infection pathology, effects on host behaviours including pest behaviours make them appealing as potential biopesticides to control their agriculturally important pest hosts. However, the efficacy of virus soldier fly virus as a viable and efficient biological control agent against solider flies remains to be established and studies investigating on host interactions, pathology and overall effects on pest population dynamics are required.

## Author and contributors

**Angelique K. Asselin:** Data curation, Formal Analysis, Investigation, Methodology, Validation, Visualization, Writing – original draft

**Kayvan Etabari:** Conceptualization, Formal Analysis, Investigation, Methodology, Validation Supervision, Writing – review & editing

**Michael J. Furlong:** Conceptualization, Funding acquisition, Project administration, Resources, Supervision, Writing – review & editing

**Karyn N. Johnson:** Methodology, Resources, Supervision, Writing – review & editing

## Acknowledgement

The authors acknowledge the facilities, and the scientific and technical assistance, of the Australian Microscopy & Microanalysis Research Facility at the Centre for Microscopy and Microanalysis, The University of Queensland.

